# Pseudogenization of the *cntQ* permease confers distinct yersinopine-metal uptake selectivity in *Yersinia* species

**DOI:** 10.1101/2025.03.14.643292

**Authors:** Clémentine Laffont, Elizabeth Pradel, Laurent Ouerdane, Amélie Dewitte, Nicolas Oswaldo Gomez, Mathilde Tribout, Catherine Brutesco, Romé Voulhoux, Ryszard Lobinski, Florent Sebbane, Pascal Arnoux

**Affiliations:** Aix Marseille Univ, CEA, CNRS, BIAM, Saint Paul-Lez-Durance, France; Univ. Lille, CNRS, Inserm, CHU Lille, Institut Pasteur de Lille, U1019 - UMR 9017 - CIIL - Center for Infection and Immunity of Lille, F-59000 Lille, France; Université de Pau et des Pays de l’Adour, e2s UPPA, CNRS, IPREM-UMR5254, Hélioparc, Pau, France; Laboratoire de Chimie Bactérienne (LCB) UMR7283, Institut Microbiologie, Bioénergies et Biotechnologie (IM2B), CNRS, Aix-Marseille Université, Marseille, France; Chair of Analytical Chemistry, Warsaw University of Technology, Poland

**Keywords:** Metal-uptake, ABC transporter, specificity, evolution, Yersinia

## Abstract

Yersinopine, a nicotianamine-like metallophore, was recently identified through biochemical analyses, but its *in vivo* production and functional role remain uncharacterized. In *Yersinia pseudotuberculosis* and its recent descendant *Yersinia pestis*, the *cnt* operon (*cntPQRLMI*) putatively encodes the biosynthesis and transport of yersinopine. In *Y. pestis*, however, two frameshift mutations disrupt *cntQ*, which encodes the predicted permease for yersinopine-metal complexes. This pseudogenization raises critical questions about the functional relevance of yersinopine in these closely related species. Here, we show that *cnt* operon expression is repressed by the zinc uptake regulator Zur and that both *Y. pestis* and *Y. pseudotuberculosis* secrete yersinopine under zinc-limited conditions. Unexpectedly, the operon mediates iron uptake in *Y. pseudotuberculosis* but supports zinc acquisition in *Y. pestis*. Moreover, targeted disruption of *cntQ* in *Y. pseudotuberculosis* shifts metal specificity from iron to zinc, mimicking the *Y. pestis* phenotype. Collectively, our results suggest that a single pseudogenization event could rewire metal uptake specificity. Our findings illustrate how evolutionary genome reduction can reshape bacterial physiology.

## Introduction

Metal uptake is fundamental for all living organisms, as metals serve as catalytic and/or structural cofactors for proteins involved in essential metabolic reactions. To efficiently acquire metals, bacteria encode diverse uptake machineries tailored to metal availability in their environmental niches (Ma et al. 2009; Chandrangsu et al. 2017). When metals are abundant or bioavailable, bacteria directly import the free metallic ions through specialized metal-uptake systems, such as ABC transporters (*e.g.,* the ZnuABC transporter responsible for zinc uptake). Under conditions of metal scarcity, metalloregulators of the FUR superfamily (such as Fur for iron or Zur for zinc) modulate gene expression to promote high-affinity acquisition pathways. These include the capture of metal-containing molecules such as haem, or the production, secretion and uptake of specialised molecules called metallophores (such as siderophores), which have a high affinity for metals, opening up new sources that would otherwise be unavailable. These high-affinity pathways are particularly important for pathogenic bacteria facing nutritional immunity, a host defense mechanism that restricts pathogen growth by sequestering essential metals or exposing microbes to toxic metal concentrations (Weinberg 1975; Hood and Skaar 2012; Zygiel and Nolan 2018; Nelson et al. 2021). Among the high-affinity metal acquisition pathways are those involving nicotianamine-like metallophores, such as staphylopine in *Staphylococcus aureus* and pseudopaline in *Pseudomonas aeruginosa* (Ghssein et al. 2016; Lhospice et al. 2017). Although initially identified for their broad metal-binding capacities, growing evidence suggests that these metallophores primarily participate in zinc acquisition in metal-chelating host environments (Grim et al. 2017; Mastropasqua et al. 2017; Lhospice et al. 2017). Nevertheless, experimental data also indicate roles in iron or nickel import (Ghssein et al. 2016; Lhospice et al. 2017; Laffont and Arnoux 2020) and staphylopine is involved in the sensitivity of *S. aureus* to copper intoxication (Hossain et al. 2023). Consistent with this versatility, the *cnt* operon governing staphylopine synthesis, secretion, and metal uptake in *S. aureus* is regulated by both zinc and iron availability (Fojcik et al. 2018). Although well characterized in *S. aureus* and *P. aeruginosa*, nicotianamine-like metallophores are widespread among pathogenic and environmental bacteria (Laffont and Arnoux 2020; Morey and Kehl-Fie 2020). Recently, biochemical analyses identified a novel nicotianamine-like metallophore, yersinopine, through *in vitro* assays using purified CntM from *Yersinia pestis*, the agent of flea-borne plague (McFarlane et al. 2018). However, its production *in vivo* and biological role remain unknown.

*Y. pestis* emerged within the past 6,000 years from *Yersinia pseudotuberculosis*, an enteropathogen that causes mild enteric infections in humans (Achtman et al. 1999). The mechanisms underlying this emergence remain partially understood but involve the acquisition and loss (through deletion or pseudogenization) of a few genes (Chain et al. 2004; Hinnebusch et al. 2016). Of interest with regard to metal homeostasis, a frameshift mutation in *ureD* abolished the nickel-dependent urease activity of *Y. pestis* (Sebbane et al. 2001). This mutation eliminated urease activity and increased flea infection incidence by reducing bacterial toxicity towards fleas (Chouikha and Hinnebusch 2014). In this context, it is noteworthy that *cntQ* (sometimes called *fiuA* or *fhuD*), encoding the predicted inner membrane importer of yersinopine-metal complexes, is disrupted in *Y. pestis* due to two frameshift mutations relative to *Y. pseudotuberculosis* (Supplementary Figure 1 for sequence alignments of the *cnt* genes; Supplementary Figures 2-4 for the AlphaFold analyses supporting the impairment of the permease; Supplementary Table 1 for the conservation of these two mutations in *Yersinia* species). Furthermore, staphylopine (an epimer of yersinopine) has been linked to urease activity in *S. aureus* (Remy et al. 2013). These observations suggest a possible link between *cntQ* disruption and loss of urease activity in *Y. pestis*. However, *cntQ* is part of the *cntPQRLMI* operon, predicted to govern yersinopine synthesis (*cntLM*) and export (*cntI*), and recovery of yersinopine-metal complexes via an ABC transporter (*cntPQR*) (Figure 1). Altogether, these observations raise the question of whether *Y. pestis* still synthesizes and secretes yersinopine, and if so, whether it contributes to metal uptake in *Y. pestis* and *Y. pseudotuberculosis*.

**Figure 1.**
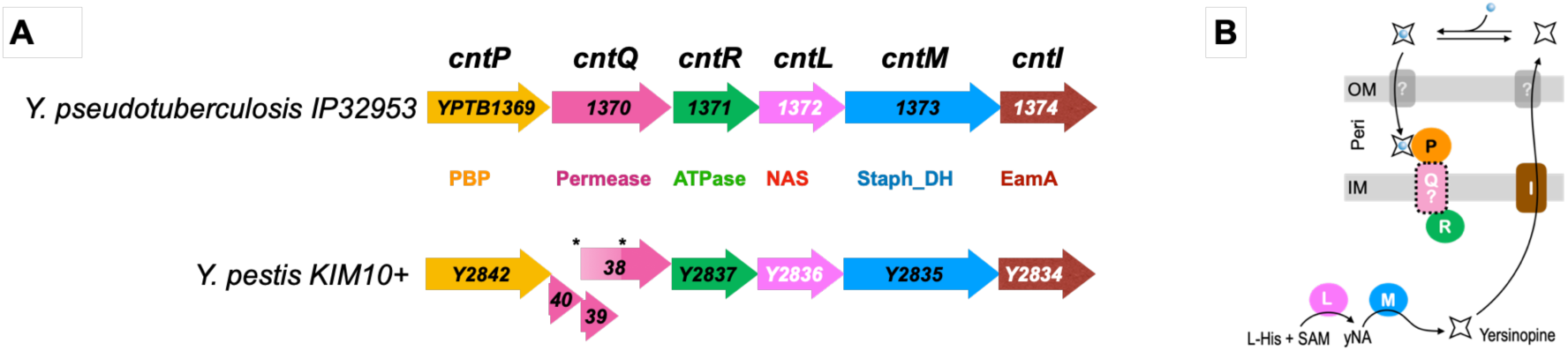
Comparison of the *cnt* operon structure in *Y. pseudotuberculosis* and *Y. pestis*. *A,* Two single-base deletions in the *Y. pestis* genome (indicated by stars) generate frameshift mutations in the *cntQ* gene. The first deletion produces a premature stop codon, leading to ORF Y2840. The second causes two additional ORFs (Y2839 and Y2838) that are frameshifted relative to *Y. pseudotuberculosis cntQ*. Notably, the second deletion restores the reading frame for the last 162 residues, resulting in partial identity between *Y. pestis* CntQ (residues 118–280) and *Y. pseudotuberculosis* CntQ (YPTB1370, residues 190–352). These three predicted ORFs are unlikely to encode a functional permease. Frameshifted region is shown in light color. Other proteins encoded by the operons share 99.3% (CntL), 99.7% (CntP), or 100% (CntR, CntM, and CntI) sequence identity between the two species. *B,* Model of yersinopine biosynthesis, export, and import of yersinopine-metal complexes in the periplasm of *Yersinia* species.

## Materials and Methods

### Bacterial strains, plasmids and culture conditions

Bacterial strains are listed in Supplementary Table 2. Cultures were grown aerobically at 28 °C with shaking in TMH minimal medium. Composition of TMH medium is listed in supplementary Table 3, and ICP-OES metal content is indicated in Supplementary Table 4. Inoculation was performed at 5.10^6^ cells/mL into 50 mL of medium in sterile 250-mL clear polystyrene bottles (Easy Grip Corning, VWR). Media were prepared with plasticware and sterilized by 0.22-μm filtration to minimize metal contamination. Growth was monitored by measuring OD_600_ over time.

### Mutant construction

*Y. pseudotuberculosis* mutants were generated by allelic exchange using suicide plasmid pDS132 derivatives in *E. coli* S17-1λpir, followed by conjugation (Simon et al. 1983) (Supplementary Table 2). *Y. pestis* mutants were constructed by Red recombination using plasmid pEP1436 as previously described (Pradel et al. 2014; Dewitte et al. 2020). In both species, antibiotic resistance cassettes were eventually removed by FLP-mediated recombination from plasmid pFLP3, which was subsequently cured by sucrose counter selection. Chromosomal *cnt*::*lacZ* fusions were generated in *Y. pseudotuberculosis* and *Y. pestis* by kanamycin selection following conjugative transfer of suicide plasmids pEP696 or pEP620. The *Y. pseudotuberculosis cntPQRLMI* operon was cloned into pEP699 by ExoCET assembly (Wang et al. 2018) and inserted into the mini*Tn*7T-Zeo2 vector pEP1046. Site-specific chromosomal integration was achieved by triparental mating with *E. coli* S17-1λpir strains carrying pEP701 and pTNS3, and verified by PCR.

### Growth curves and doubling time

Overnight cultures were grown aerobically at 28°C in TMH medium using polycarbonate Erlenmeyer flasks. Three independent biological replicates were inoculated at an initial OD_600_ of 0.05 into fresh TMH medium (200 μL) supplemented with various concentrations of EDTA (0–800 µM) in 96-well flat-bottom plates (Corning Costar 3596). Growth was monitored every 15 min for 24 hours at 28°C using a Tecan Spark Microplate Reader under continuous shaking. OD_600_ values were corrected for background absorbance using non-inoculated media. Growth curves were analyzed in RStudio, and doubling times were calculated using the Growthcurver package (Sprouffske and Wagner 2016).

### Genome analysis, protein cloning, expression and purification

The synteny and sequence conservation of *cnt* genes across *Y. pestis* and *Y. pseudotuberculosis* strains were analyzed using the MaGe MicroScope platform (Vallenet et al. 2020). The *cntL* and *cntM* genes (*yptb1372–1373*) were amplified from *Y. pseudotuberculosis* 2777 genomic DNA using Q5® High-Fidelity DNA polymerase (New England BioLabs) and cloned into the pET-SUMO vector (Champion™ pET SUMO Protein Expression System, Invitrogen). C-terminal His-tagged proteins were expressed *in E. coli* BL21(DE3) carrying pRARE and purified by nickel affinity chromatography (HisTrap column, GE Healthcare) according to the manufacturer’s instructions. Eluted proteins were dialyzed into imidazole-free buffer (20 mM Na₂HPO₄, 300 mM NaCl, 10% glycerol, pH 7.5) and analyzed by SDS-PAGE (NuPAGE 10% Bis-Tris gels, Invitrogen).

### Detection of yersinopine by TLC assay

TLC assays were performed as previously described for staphylopine, pseudopaline, and bacillopaline detection (Higuchi et al. 1999; Ghssein et al. 2016; Lhospice et al. 2017; Laffont et al. 2019). Purified CntL and CntM proteins (2.5 μM each) were incubated with carboxyl-[^14^C]-labeled SAM (2.5 μM), L- or D-histidine (1 mM), NADPH or NADH (10 μM), and pyruvate or α-ketoglutarate (1 mM) in 50 mM HEPES buffer (pH 9) containing 1 mM DTT and 1 mM EDTA. Reactions (100 μL) were carried out for 30 min at 28°C and stopped by adding ethanol to 50% (v/v). Samples (10 μL) were spotted onto HPTLC Silica Gel 60 plates (Merck) and developed with a phenol:n-butanol:formate:water (12:3:2:3, v/v) solvent system. After drying, plates were exposed to [^14^C]-sensitive imaging plates (GE Healthcare) for 24 h and scanned with a Typhoon FLA 7000 phosphorimager.

### Sample preparation and detection of yersinopine by HILIC/ESI-MS

For *in vitro* assays, purified CntL and CntM (2.5 μM each) were incubated with SAM (2.5 μM), L-histidine (1 mM), NADPH or NADH (10 μM), and pyruvate or α-ketoglutarate (1 mM) in 50 mM HEPES buffer (pH 9) supplemented with 1 mM DTT and 1 mM EDTA. Reactions were carried out at 28°C for 30 min, then frozen at −80°C until analysis. For extracellular samples, supernatants of *Y. pestis* and *Y. pseudotuberculosis* cultures grown for 24 h at 28°C in TMH medium were collected after centrifugation (5,000 × g, 20 min, 4°C), filtered (0.22 µm), and stored at −80°C. Samples were analyzed by hydrophilic interaction liquid chromatography coupled with electrospray ionization mass spectrometry (HILIC/ESI-MS). Separations were performed on a TSKgel Amide-80 column (Tosoh Biosciences) using a linear gradient of acetonitrile and 10 mM ammonium formate (pH 5.5) at a flow rate of 50 µL/min. An Agilent 1100 HPLC system coupled to an LTQ Orbitrap Velos mass spectrometer (Thermo Fisher Scientific) was used. Prior to injection, samples were diluted 1:2 with acetonitrile, centrifuged, and injected (7 µl). MS spectra were recalibrated offline using internal standards.

### Analysis of metal concentration by ICP-MS

*Y. pestis* and *Y. pseudotuberculosis* grown in TMH medium under aerobic conditions at 28°C for 24 h were centrifuged (4,000 g, 20 min, 4°C). Bacterial pellets were washed with 20 mM ammonium acetate, dried at 50°C for 48 h, and digested with a mixture of 70% HNO₃ and 30% H₂O₂. After heating at 80°C for 3 h, samples were diluted to 2 mL with water and analyzed by inductively coupled plasma mass spectrometry (ICP-MS, Agilent 7500cs) operating in hydrogen collision gas mode.

### Statistical analysis

ICP-MS measurements were performed on two or three independent biological triplicates, collected on different days. For each experiment, metal content was normalized to the corresponding wild-type strain. Statistical analyses were conducted in RStudio (version 4.1.1). Differences between mutant and wild-type strains were assessed using Welch’s t-test after normality verification. p-values were adjusted for multiple testing using the Bonferroni method.

## Results

### Biochemical evidence that yersinopine is produced by Y. pseudotuberculosis

Yersinopine production was previously inferred through biochemical assays of the CntM enzyme from *Y. pestis* under steady-state kinetic conditions (McFarlane et al. 2018), but the full biosynthetic pathway remained incomplete because CntL was not purified. Here, we completed the pathway by characterizing purified recombinant CntL and CntM from *Y. pseudotuberculosis* (Supplementary Figure S5) using multiple biochemical assays. First, we used a ^14^C-labelled SAM substrate assay to trace the aminobutyrate moiety of SAM in the presence of L-histidine or D-histidine, followed by separation on a TLC plate. This approach demonstrated that CntL indeed specifically uses SAM and L-histidine to generate the yNA intermediate (Figure 2(A)). Building on this result, co-incubation of CntL and CntM with SAM, L-histidine, α-ketoacids, and NAD(P)H showed that CntM utilizes NADPH to reductively condense yNA with pyruvate, yielding yersinopine (Figure 2(B)). HILIC/ESI-MS analysis further validated the use of L-histidine (Figure 2(C)) and pyruvate (Figure 2(D)) by CntL and CntM, respectively. Collectively, our results demonstrate that CntL and CntM catalyze the formation of yersinopine according to the reaction pathway shown in Figure 2E. Given the 99.3% and 100% sequence identity of CntL and CntM between *Y. pseudotuberculosis* and *Y. pestis*, it is highly likely that *Y. pestis* produces an identical metallophore, namely yersinopine.

**Figure 2.**
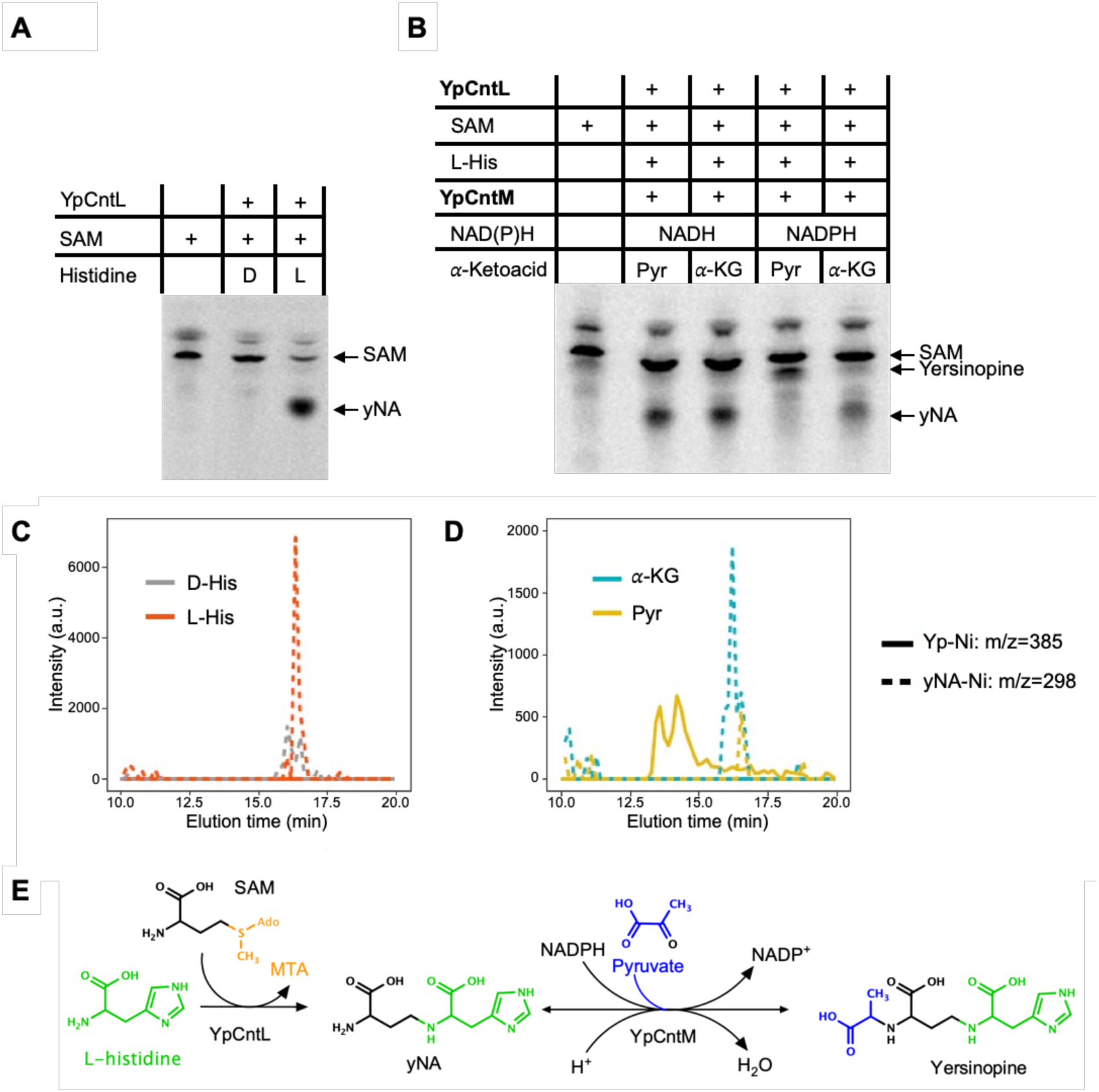
*In vitro* selectivity of CntL and CntM. *A,* Radiogram showing the activity of CntL after incubation with ^14^C-SAM and either L- or D-histidine. *B,* Radiogram after co-incubation of CntL (with ^14^C-SAM and L-histidine) and CntM with pyruvate or α-ketoglutarate (α-KG) and NAD(P)H. *C,* HILIC/ESI-MS detection of the yNA-nickel complex (dashed line) after incubation of CntL with SAM and either L- or D-histidine. *D*, HILIC/ESI-MS detection of yNA-nickel (dashed line) or yersinopine-nickel (solid line) complexes after co-incubation of CntL (with SAM and L-histidine) and CntM (with NADPH and either pyruvate or α-KG). *E,* Schematic representation of the overall yersinopine biosynthetic pathway.

### Yersinopine is produced and exported by both Y. pseudotuberculosis and Y. pestis

After establishing the roles of CntL and CntM in yersinopine biosynthesis, and inferring that CntI mediates its export across the inner membrane (Lhospice et al. 2017; McFarlane et al. 2018; Laffont et al. 2019), we next investigated whether yersinopine is produced and secreted by both *Y. pestis* and *Y. pseudotuberculosis*. To this end, we quantified yersinopine in the extracellular fractions of both bacteria grown in TMH medium using HILIC coupled with ESI-MS (Figure 3). Yersinopine was detected complexed with multiple divalent metals, predominantly zinc, and to a lesser extent copper, nickel, and cobalt. This distribution might be a proxy for the higher affinity of yersinopine for zinc under these conditions. Iron(II)-yersinopine complexes were not observed, likely due to oxidation under aerobic conditions. Metal-yersinopine complexes were detected in the supernatants of both species, although the abundance was lower in *Y. pestis*, suggesting a reduced extracellular concentration compared to *Y. pseudotuberculosis*. Collectively, these results demonstrate that both species produce and secrete yersinopine. Moreover, they reveal that the pseudogenization of *cntQ* in *Y. pestis* does not impair the operon’s ability to govern yersinopine biosynthesis and secretion.

**Figure 3.**
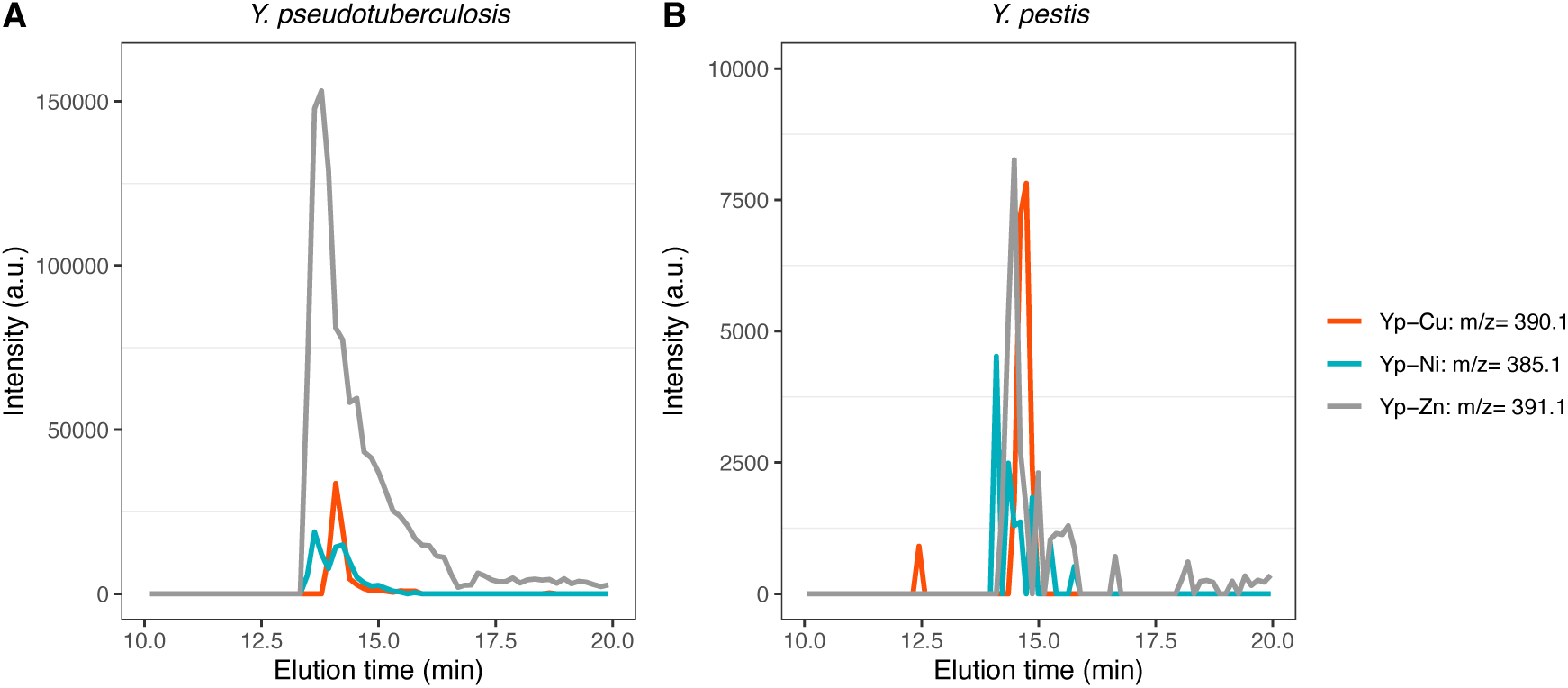
*In vivo* production and secretion of yersinopine by *Y. pseudotuberculosis* and *Y. pestis*. HILIC/ESI-MS detection of divalent yersinopine-metal complexes in culture supernatants of *Y. pseudotuberculosis* (A) and *Y. pestis* (B).

### The cnt operon is repressed by Zur in both Y. pseudotuberculosis and Y. pestis

Previous studies of the Zur regulon in *Y. pestis* (Li et al. 2009; Desrosiers et al. 2010) and *Y. pseudotuberculosis* (Gu et al. 2024) revealed strong upregulation of *cntP* in Zur-deficient strains (227-fold in *Y. pestis*, 9-fold in *Y. pseudotuberculosis*), along with induction of downstream operon genes (ranging from 45-fold to 6-fold depending on the gene). However, at that time, the function of the *cnt* system remained unclear. Although Zur was also implicated in zinc homeostasis and oxidative stress defense in *Y. pseudotuberculosis* (Wang et al. 2016), the role of the *cnt* system itself was not addressed. Given that both *Y. pseudotuberculosis* and *Y. pestis* produce yersinopine, we next investigated whether the *cnt* operon is subject to Zur regulation. To test this, we inserted a *lacZ* transcriptional reporter downstream of the *cnt* promoter in Zur^+^ and Zur^−^ strains of both species. As expected, promoter activity was repressed in Zur^+^ strains and derepressed in Zur^−^mutants grown in rich medium (Supplementary Figure 6). These findings confirm that Zur represses the *cnt* operon in both *Y. pestis* and *Y. pseudotuberculosis* in zinc-repleted conditions.

### Yersinopine mediates distinctive strategies of metal uptake in Y. pseudotuberculosis and Y. pestis

Since the *cnt* operon is repressed by Zur and involved in yersinopine production and secretion, we next examined the impact of *cnt* deletions on metal accumulation. To explore the role of the *cnt* operon, we generated mutants in both species, including a strain lacking *cntQ* with a polar effect, abolishing yersinopine production (Δ*cntQ(RLMI)*), and a strain lacking *cntLMI* (Δ*cntLMI*; see Methods). In *Y. pseudotuberculosis*, we also generated a non-polar Δ*cntQ* mutant to mimic the genetic configuration of *Y. pestis*. When TMH medium was supplemented with EDTA (a metal chelator commonly used to trigger metal deficiency) the growth rate of the *Y. pseudotuberculosis* wild-type strain increased, whereas *cnt* mutants exhibited reduced growth (Supplementary Figure 7). This suggests that the *cnt* operon provides a growth advantage under these metal-limited conditions. However, in order to compare wild-type and mutant strains across both species, we grew them in TMH medium without EDTA (conditions in which we established yersinopine secretion by both species) and measured intracellular metal content by inductively coupled plasma mass spectrometry (ICP-MS).

In *Y. pseudotuberculosis*, loss of the yersinopine biosynthetic genes (Δ*cntQ(RLMI)* and Δ*cntLMI*) led to a slight but significant reduction in intracellular iron content (−27.6%, *p* = 3.7×10^−5^ and −31.1%, *p* = 4.5×10^−3^, respectively), without affecting zinc or other metals (Figure 4(A)). Complementation restored iron to wild-type levels. By contrast, in *Y. pestis,* iron content was unchanged, but zinc levels decreased significantly in both Δ*cntQ(RLMI)* and Δ*cntLMI* mutants (−47.3%, *p* = 6.2×10^−3^ and −50.9%, *p* = 1.9×10^−2^), while cobalt levels rose substantially (+176.4%, p=2.8×10^−6^ and +182.2%) (Figure 4(B)).

**Figure 4.**
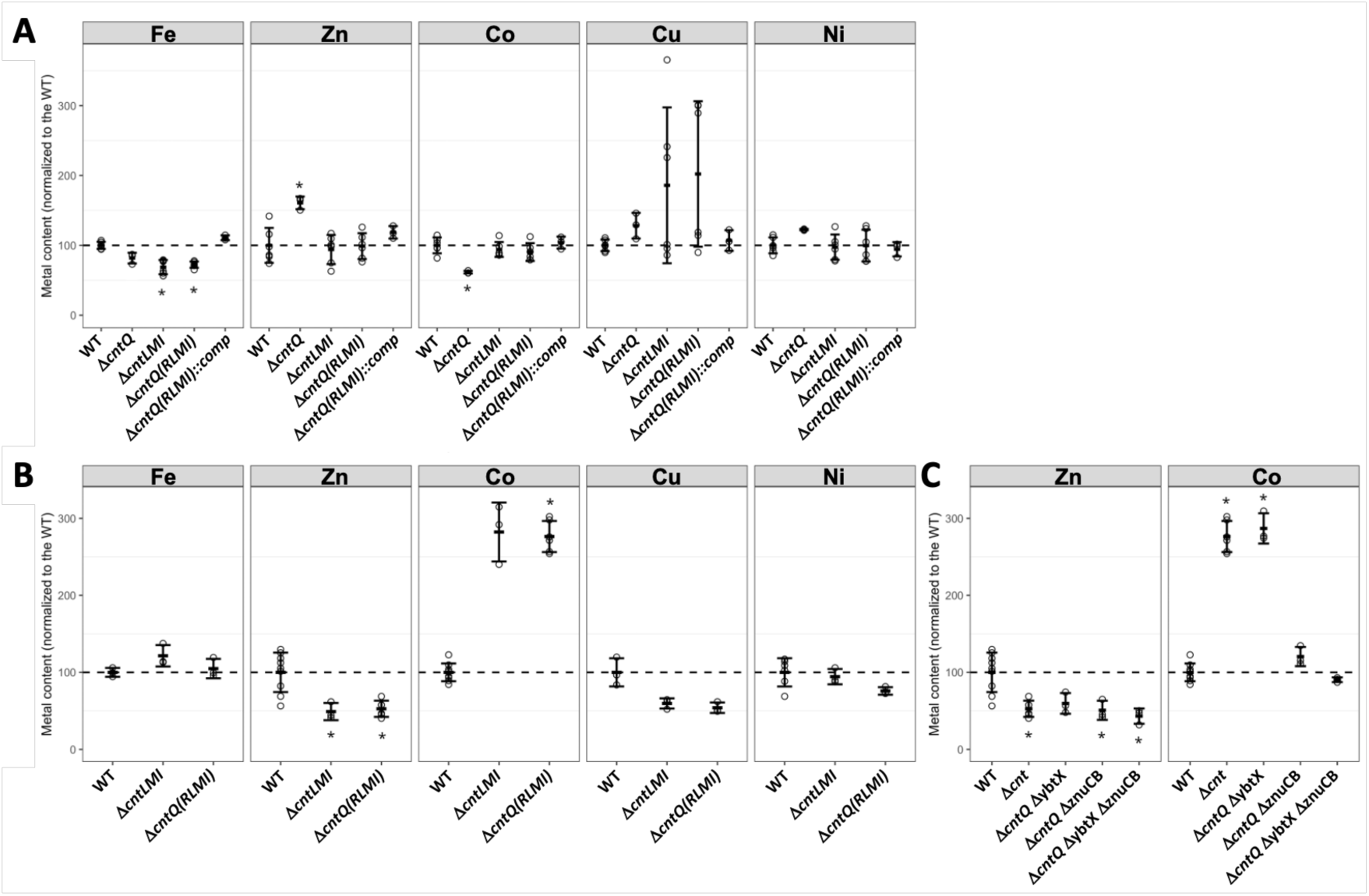
Intracellular metal content in *Y. pseudotuberculosis* and *Y. pestis* strains. *A,* ICP-MS quantification of intracellular metals in *Y. pseudotuberculosis* wild-type (WT) and *cnt* mutants. *B,* ICP-MS quantification of intracellular metals in *Y. pestis* WT and *cnt* mutants. *C,* Intracellular zinc and cobalt levels in *Y. pestis* WT and strains lacking the Cnt system alone or combined with deletions of the yersiniabactin transporter (YbtX) and/or the ZnuABC zinc importer. Data are from 1 to 3 independent triplicates (total n= 3-9). Asterisks (*) indicate significant differences relative to WT (Welch’s t-test with Bonferroni correction; p < 0.05).

To further explore zinc and cobalt uptake, we analyzed *Y. pestis Δcnt* mutants additionally lacking either the putative yersiniabactin importer YbtX (which mediates yersiniabactin-zinc uptake) (Price et al. 2021), the Znu zinc uptake system, or both (Figure 4(C)). Compared to the yersinopine-deficient mutant, intracellular zinc concentrations did not decrease further in the double and triple mutants, indicating that unidentified zinc uptake mechanisms remain active. In contrast, cobalt accumulation returned to wild-type values in the Δ*cnt* Δ*znu* and Δ*cnt* Δ*ybtX* Δ*znu* mutants, but remained elevated in mutants lacking only *cnt* or both *cnt* and *ybtX*. These findings suggest that in the absence of yersinopine, cobalt can be imported through Znu, whereas the presence of yersinopine may inhibit cobalt uptake via Znu in *Y. pestis*.

### Deletion of cntQ in Y. pseudotuberculosis mimics the evolutionary shift in metal-uptake specificity

Our findings suggest that during the emergence of *Y. pestis* from *Y. pseudotuberculosis*, pseudogenization of *cntQ* reshaped metal-uptake specificity from iron to zinc. To directly test this hypothesis, we deleted *cntQ* in *Y. pseudotuberculosis*. Strikingly, the reduction in intracellular iron observed in yersinopine-deficient strains was no longer detectable. Instead, *cntQ* deletion triggered a significant rise in zinc (+60.8%, *p* = 2.5×10^−2^) and a concomitant fall in cobalt (−38.4%, *p* = 5.8×10^−3^) (Figure 4(A)). This shift recapitulates the metal-uptake profile of wild-type *Y. pestis*, where the Cnt system promotes zinc acquisition while restricting cobalt accumulation. Thus, production of yersinopine combined with a defective yersinopine-metal complex inner membrane importer rewires metal transport toward zinc uptake and limits cobalt entry via the Znu system, closely mirroring the transport dynamics observed in *Y. pestis*.

## Discussion

A functional yersinopine-mediated metal uptake system (*cntPQRLMI*) was predicted in *Y. pseudotuberculosis*, but the presence of a frameshift mutation in the *cntQ* permease gene suggested that this system was non-functional in *Y. pestis* (Forman et al. 2010). Here, we demonstrate that yersinopine is produced and secreted by both *Y. pseudotuberculosis* and *Y. pestis* when grown in the same low-zinc medium, indicating that production and export are preserved despite the inactivation of *cntQ* in *Y. pestis*. These findings suggest that the *cnt* operon retains some function despite the pseudogenization of *cntQ*. This is further supported by transcriptomic data showing that the *Y. pestis cnt* operon is controlled by the Zur repressor (Li et al. 2009) and is strongly induced during pneumonic infection (Perry et al. 2015). Therefore, an important question that arises is the role of yersinopine in the context of either a fully functional Cnt system (*Y. pseudotuberculosis*) or a system lacking the ABC importer permease (*Y. pestis*).

Our data suggest that the Cnt system is not involved in nickel uptake in *Y. pestis* and *Y. pseudotuberculosis*, in contrast to findings in *P. aeruginosa* (Mastropasqua et al. 2017; Lhospice et al. 2017) and *S. aureus* (Remy et al. 2013). Unexpectedly, our results indicate that the presence or absence of the CntQ permease influences metal-uptake specificity in *Yersinia*, shifting from an iron uptake system when CntQ is present (*Y. pseudotuberculosis*) to a zinc uptake system when CntQ is absent (*Y. pestis*). The main limitation of our study is that we do not differentiate between compartmentalization of metal accumulation, *i.e.* this could occur in the periplasm as well as in the cytoplasm. To our knowledge, this represents a unique case where pseudogenization of a permease within an ABC transporter leads to a shift in substrate specificity, even if metals may accumulate in different compartments of the cell. Interestingly, a frameshift mutation in the *iucA* gene, which encodes a key enzyme in aerobactin siderophore biosynthesis, has also been identified in *Y. pestis*. However, the bacterium remains capable of utilizing aerobactin-iron complexes for iron uptake, presumably through piracy of aerobactin produced by other bacteria in its biotope (Forman et al. 2007). The extent to which our finding that the yersinopine system of *Y. pestis* is involved in zinc homeostasis *in vitro* translates to host-associated or other environmentally relevant conditions remains to be evaluated.

Functional studies of the Znu system in *Y. pestis* indicated that it is not required for virulence, suggesting the existence of additional zinc-uptake mechanisms (Desrosiers et al. 2010). Among these, yersiniabactin-mediated zinc acquisition was later recognized (Price et al. 2021). Our findings further expand this understanding by revealing that the *cnt* operon also plays a critical role in zinc uptake. While the entire operon appears essential, the route of zinc entry in the cell is unknown and the specific role of the periplasmic protein CntP remains uncertain. It may either be rerouted to an alternative transporter or act indirectly, for instance by chelating a yersinopine-metal complex. Consistent with this, the very likely impairment of CntQ in *Y. pestis* supports the hypothesis that yersinopine functions independently of CntP or CntP-R. In this context, analyzing mutants lacking *cntPQR* in both *Y. pestis* and *Y. pseudotuberculosis* would be particularly informative to clarify CntP’s role in metal acquisition.

The increased intracellular cobalt accumulation in *Y. pestis* upon inactivation of the *cnt* operon was unexpected. However, we found that this increase was correlated with the presence of a functional Znu system, as the absence of both Cnt and Znu systems restored intracellular cobalt levels to wild-type values. This interplay between the Znu and Cnt systems could be explained by the ability of yersinopine to form a complex with cobalt in the periplasmic space, thereby preventing cobalt entry into the cytoplasm via the Znu system. A connection between Znu and cobalt transport has been previously described in *P. aeruginosa* (Pederick et al. 2015; Mastropasqua et al. 2017), where cobalt uptake was found to increase in a mutant lacking the ZnuABC transporter, suggesting a zinc-regulated cobalt uptake mechanism. Although the mechanisms differ, both our findings and prior studies underscore the frequent cross-talk between zinc and cobalt uptake systems. A model summarizing zinc and cobalt import via the Cnt and Znu systems in both *Y. pseudotuberculosis* and *Y. pestis* is presented in Figure 5. This model also illustrates how small genetic changes can significantly alter bacterial physiology, potentially influencing pathogen evolution and adaptation.

**Figure 5.**
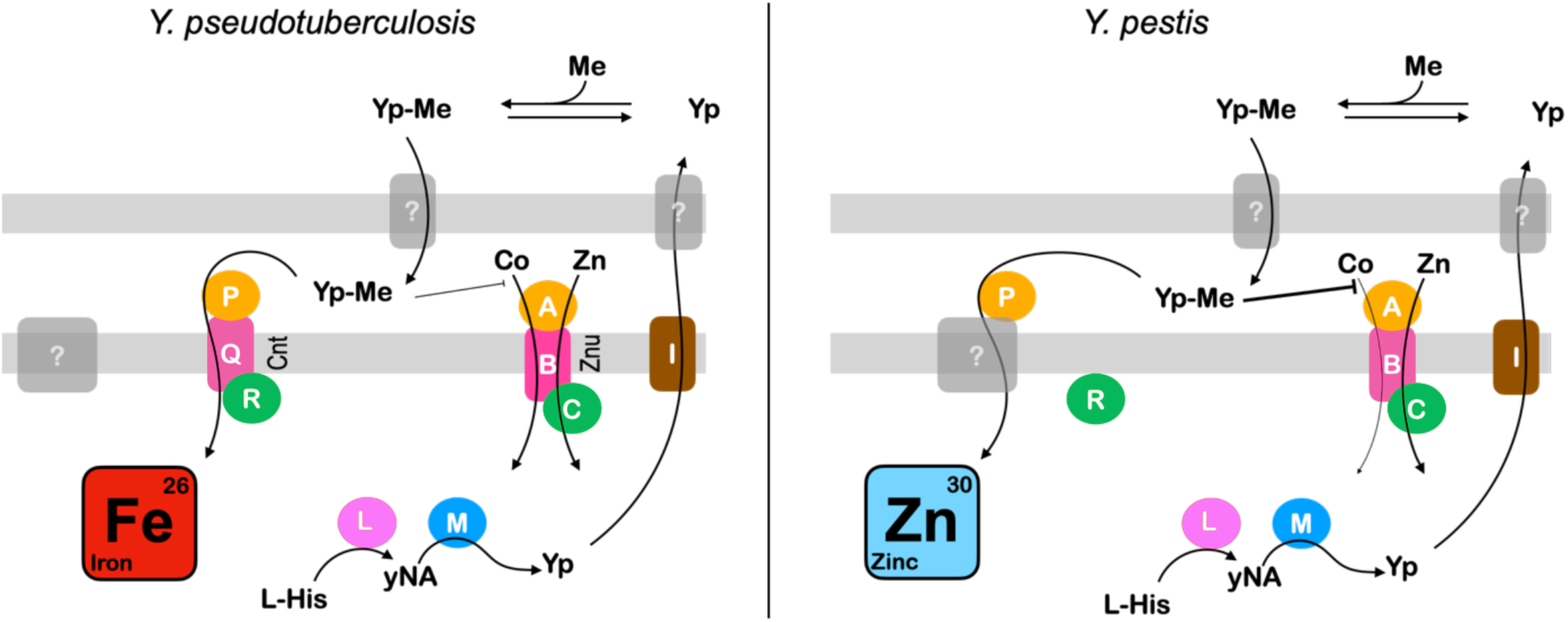
Comparative model of metal uptake through the Cnt and Znu systems in *Y. pseudotuberculosis* and *Y. pestis*. In *Y. pseudotuberculosis*, yersinopine-metal complexes in the periplasm facilitate iron uptake, presumably via the CntPQR ABC transporter encoded by the *cnt* operon. In *Y. pestis*, the absence of the CntQ permease, due to a frameshift mutation, alters metal homeostasis. Zinc import via the Cnt system appears to occur through rerouting of the yersinopine-zinc complex to an unknown inner membrane transporter (indicated by “?”), potentially involving CntP in the periplasm. In both *Yersinia* species, two additional unknown steps at the outer-membrane are proposed, including the export of yersinopine to the extracellular space and the import of yersinopine-metal complexes into the periplasm.

## Statements and Declarations

The authors declare no competing interests.

## Acknowledgements

We thank the Agence Nationale de la Recherche for initial support [grants ANR-14-CE09-0007-02 and ANR-20-CE11-0018], the French ministry of research for a PhD grant awarded to CL. FS was partly supported by the ERC Synergy project Synergy-Plague (101118880). We would like to thank Pierre Lê-Bury and Javier Pizarro-Cerda (Institut Pasteur) as well as members of both the Arnoux and Sebbane teams for their support and discussions. Richard C. Allen and Ikram Hammar are acknowledged respectively for his advice on statistical analysis and her initial illustration of the model. S. Preveral and the Ionotec plateform are recognized for their determination of metal content in TMH medium. This article has been deposited to biorxiv (https://doi.org/10.1101/2025.03.14.643292)

## Supplemental material

**Supplementary Figure 1.**
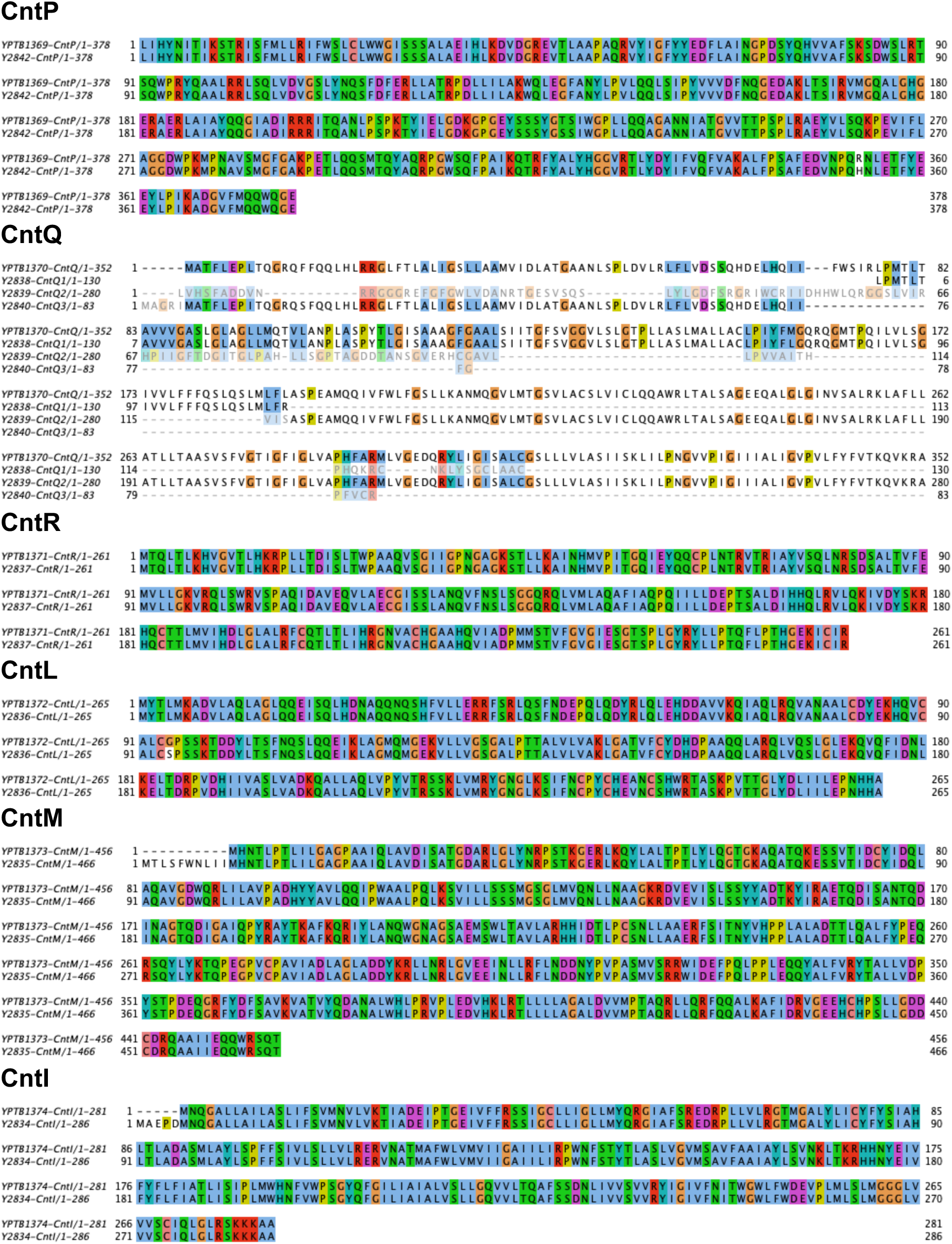
Sequence conservation between the different predicted cnt genes in *Y. pestis* and *Y. pseudotuberculosis*.

**Supplementary Figure 2.**
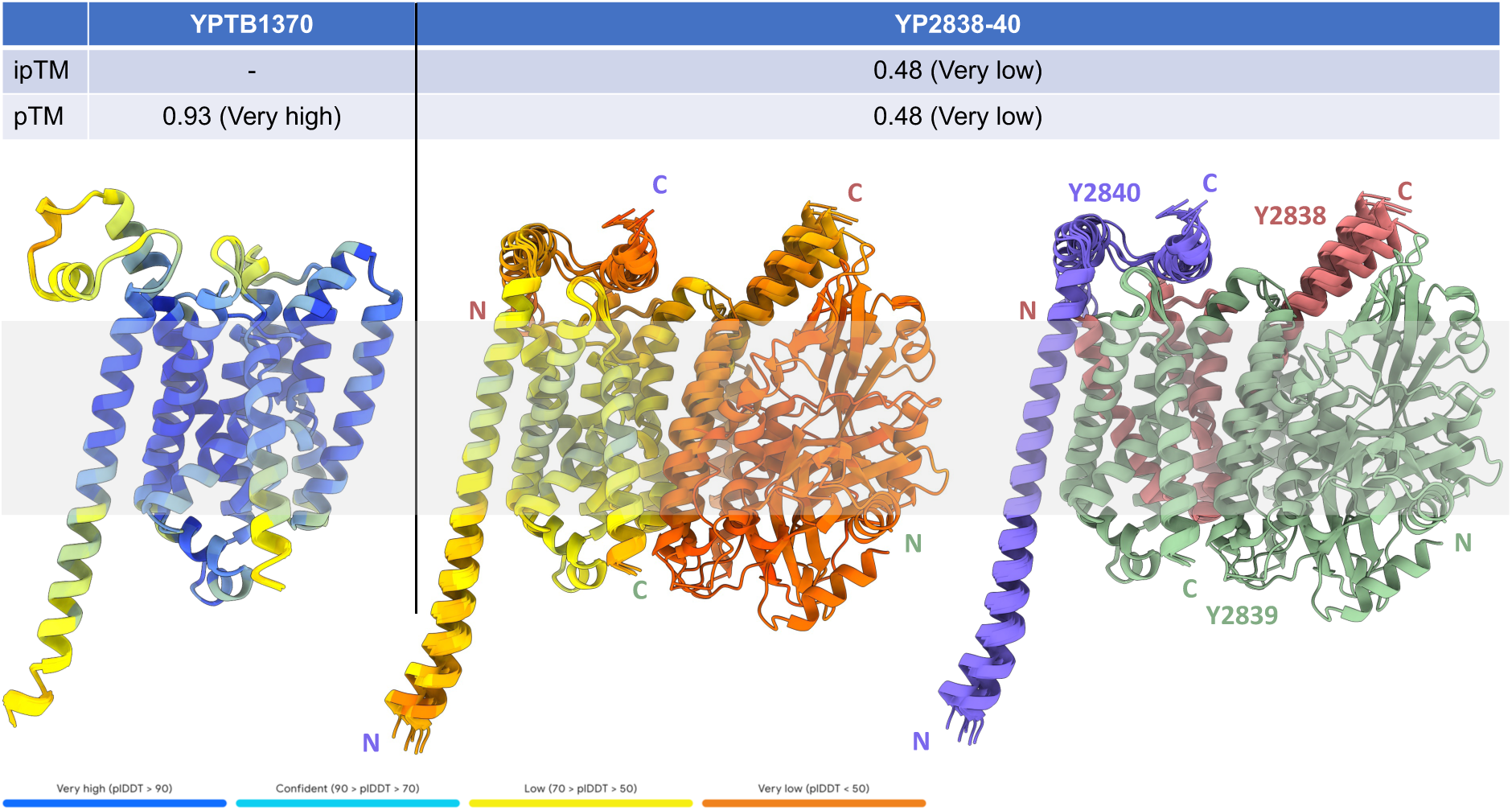
AlphaFold3 models of a CntQ monomer from *Y. pseudotuberculosis* (YPTB1370; left) and *Y. pestis* (YP2838-40; right). The models are coloured according to the quality of the prediction, except for the right panel, which shows the position of the three peptides and the location of their N- and C-terminus. The approximate position of the membrane is shown in grey.

**Supplementary Figure 3.**
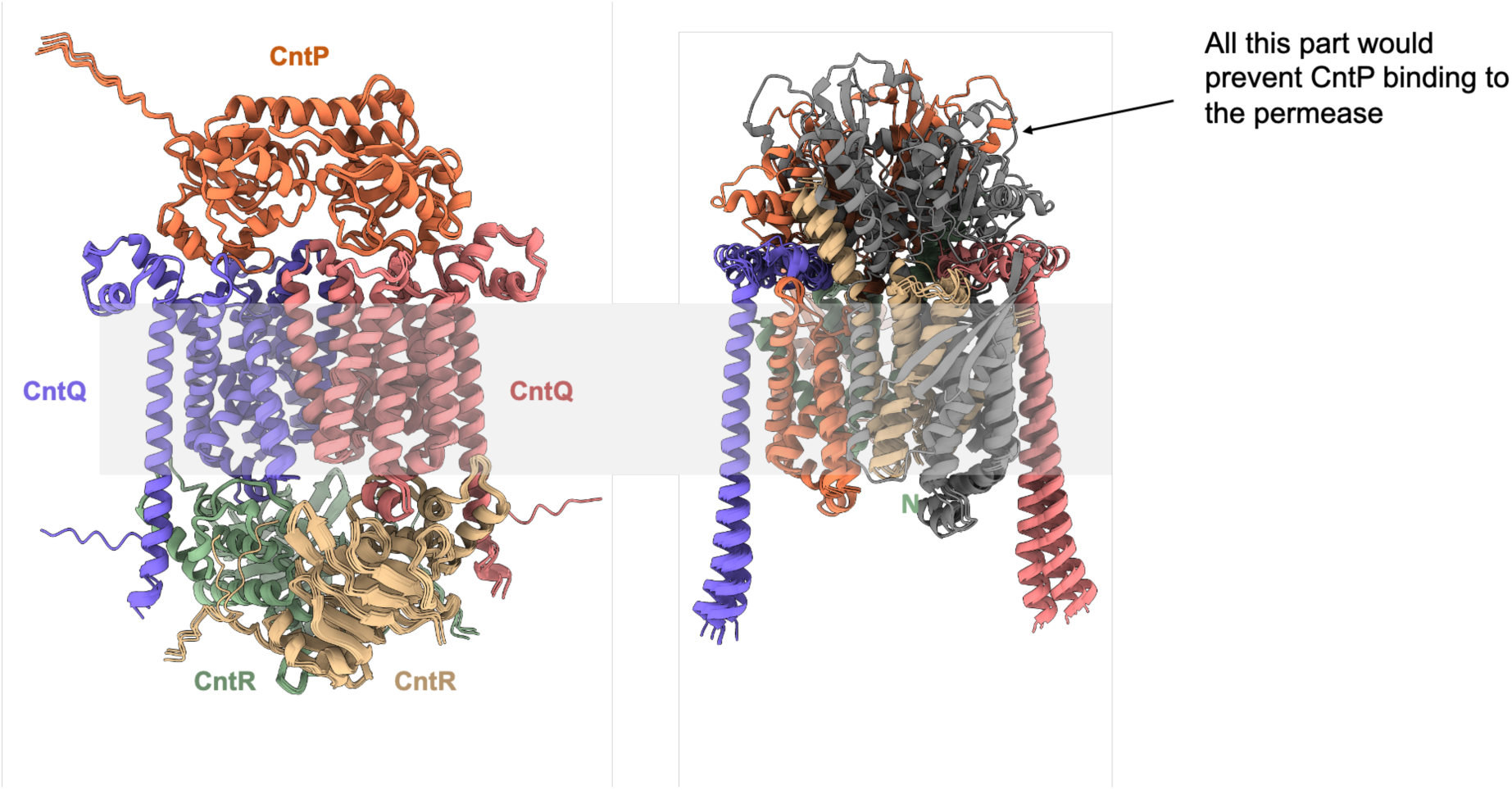
AlphaFold3 models of the CntPQ_2_R_2_ from *Y. pseudotuberculosis* (left ; ipTM = 0.81 pTM = 0.84) and the CntQ_2_ from *Y. pestis* (right ; ipTM = 0.61 pTM = 0.61). Although there would be enough space for interaction with the ATPase CntR in the cytoplasm, the space used for recognition of CntP in the periplasm is occupied by additional peptide parts that have no counterpart in *Y. pseudotuberculosis*, therefore leading to our central hypothesis that the three peptides produced by Y. pestis (Y2838-40), even if produced, are likely non-functional.

**Supplementary Figure 4.**
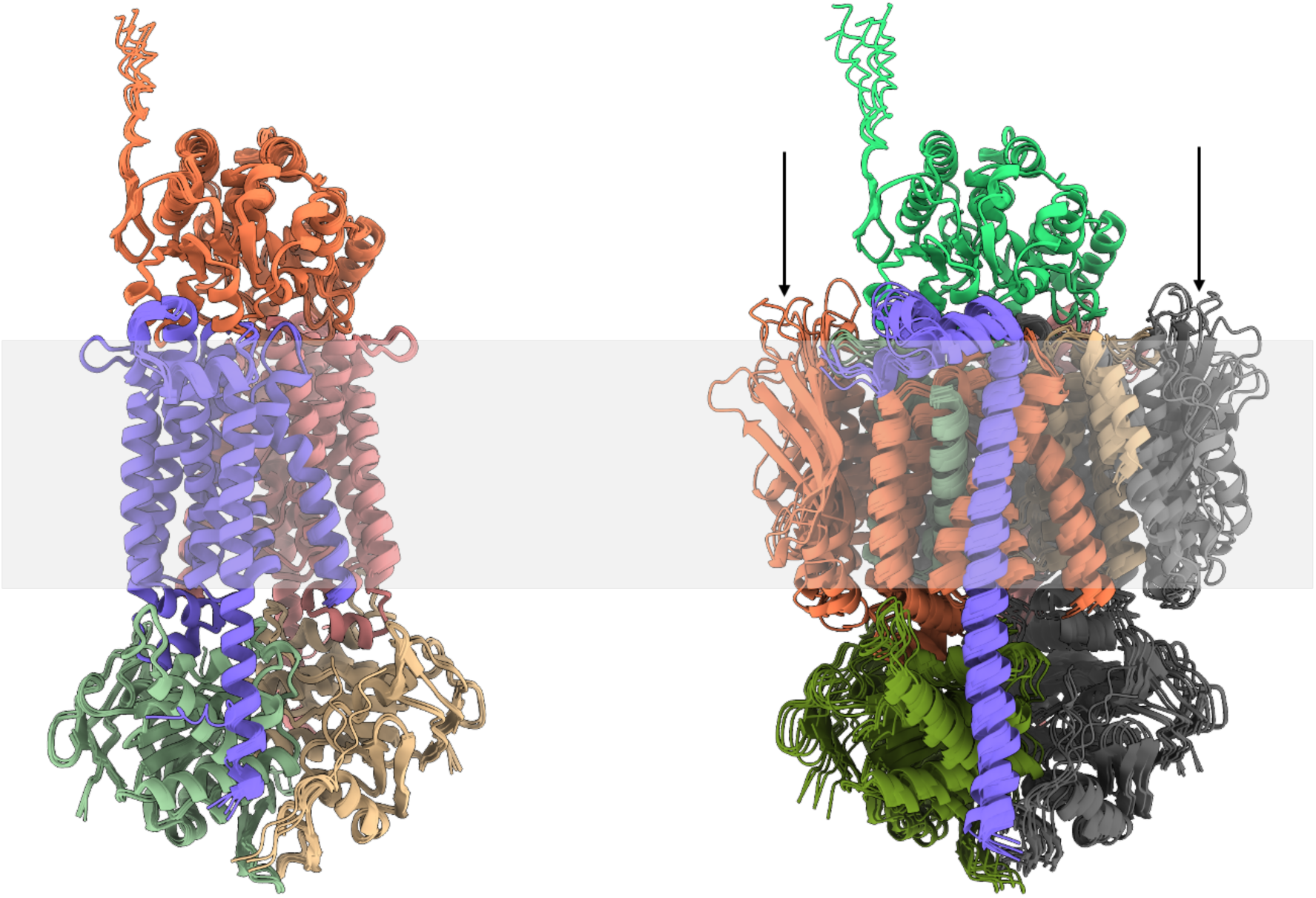
AlphaFold3 models of the CntPQ_2_R_2_ from *Y. pseudotuberculosis* (left; ipTM = 0.81 pTM = 0.84) and the equivalent from *Y. pestis* (right; ipTM = 0.65 pTM = 0.65). With regard to *Y. pestis*, the modelling with CntP indicates that the parts that have no equivalent in *Y. pseudotuberculosis* now folds in the membrane (arrows), most probably because the model forces CntP binding. However, if stable, this part should fold in the periplamic space, which should therefore prevent either membrane insertion of CntQ or CntP binding to CntQ, anyway likely impairing its function.

**Supplementary Figure 5:**
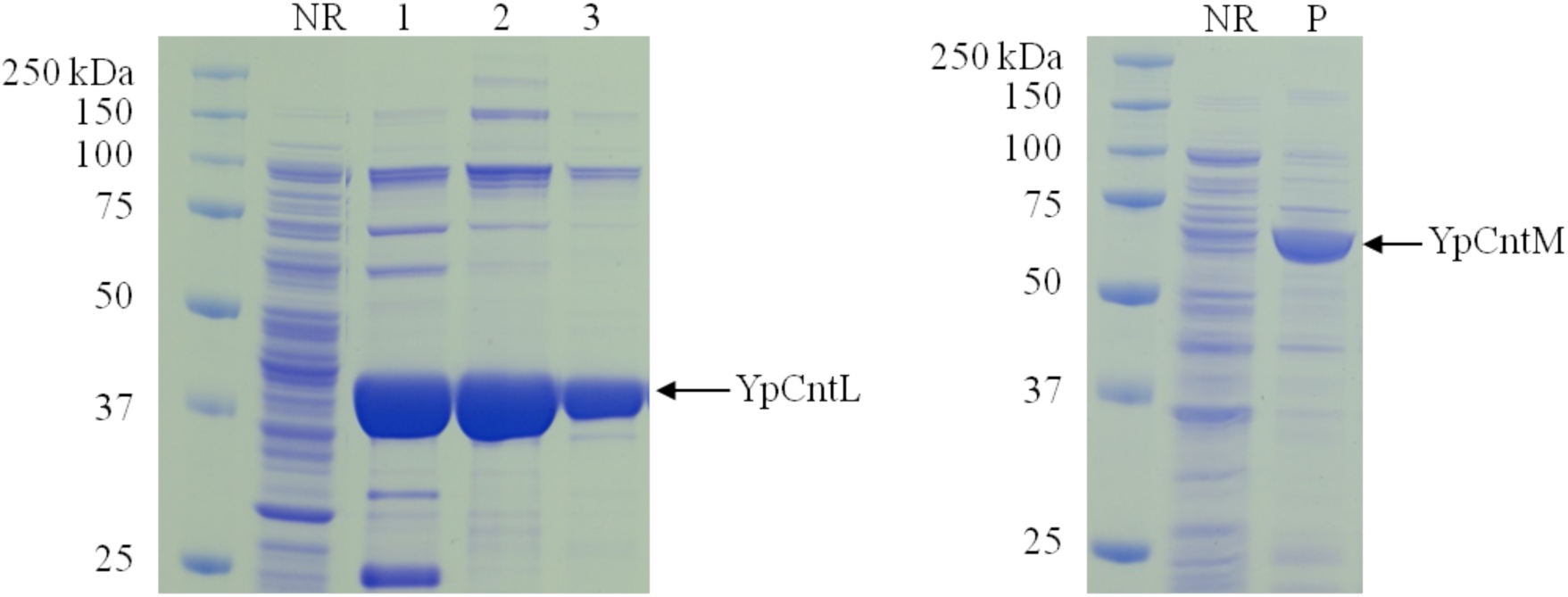
SDS-PAGE gels illustrating the purifications of YpCntL (left) and YpCntM (right) using HisTrap columns. NR corresponds to the flow-through fraction, 1, 2 and 3 to the different eluted fractions, and P to the eluted fraction of CntM.

**Supplementary Figure 6.**
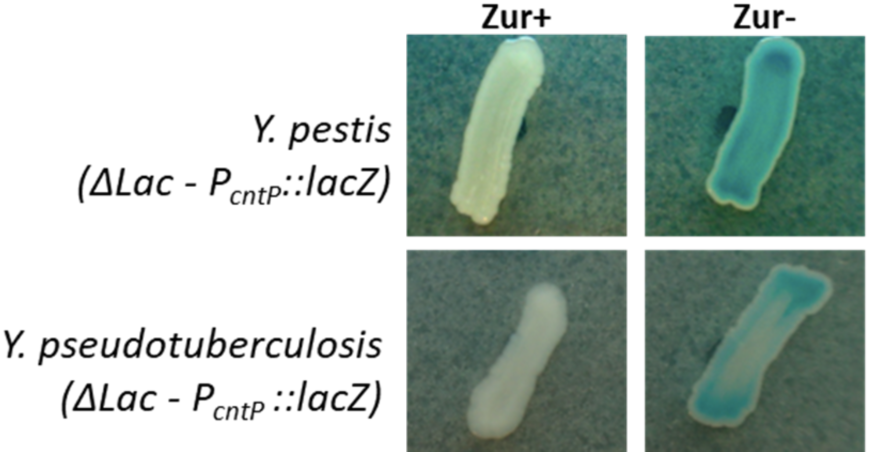
Zur-dependent repression of the *cnt* operon in *Y. pestis* and *Y. pseudotuberculosis*. *Y. pestis* and *Y. pseudotuberculosis* Δ*lacZ* strains, either expressing (+) or lacking (*-*) Zur, carried a chromosomal fusion between the *cnt* operon promoter (P*_cntP_*) and the *E. coli lacZ* reporter gene. Promoter activity was assessed by β-galactosidase activity, visualized as blue coloration on LB agar plates supplemented with X-gal.

**Supplementary Figure 7:**
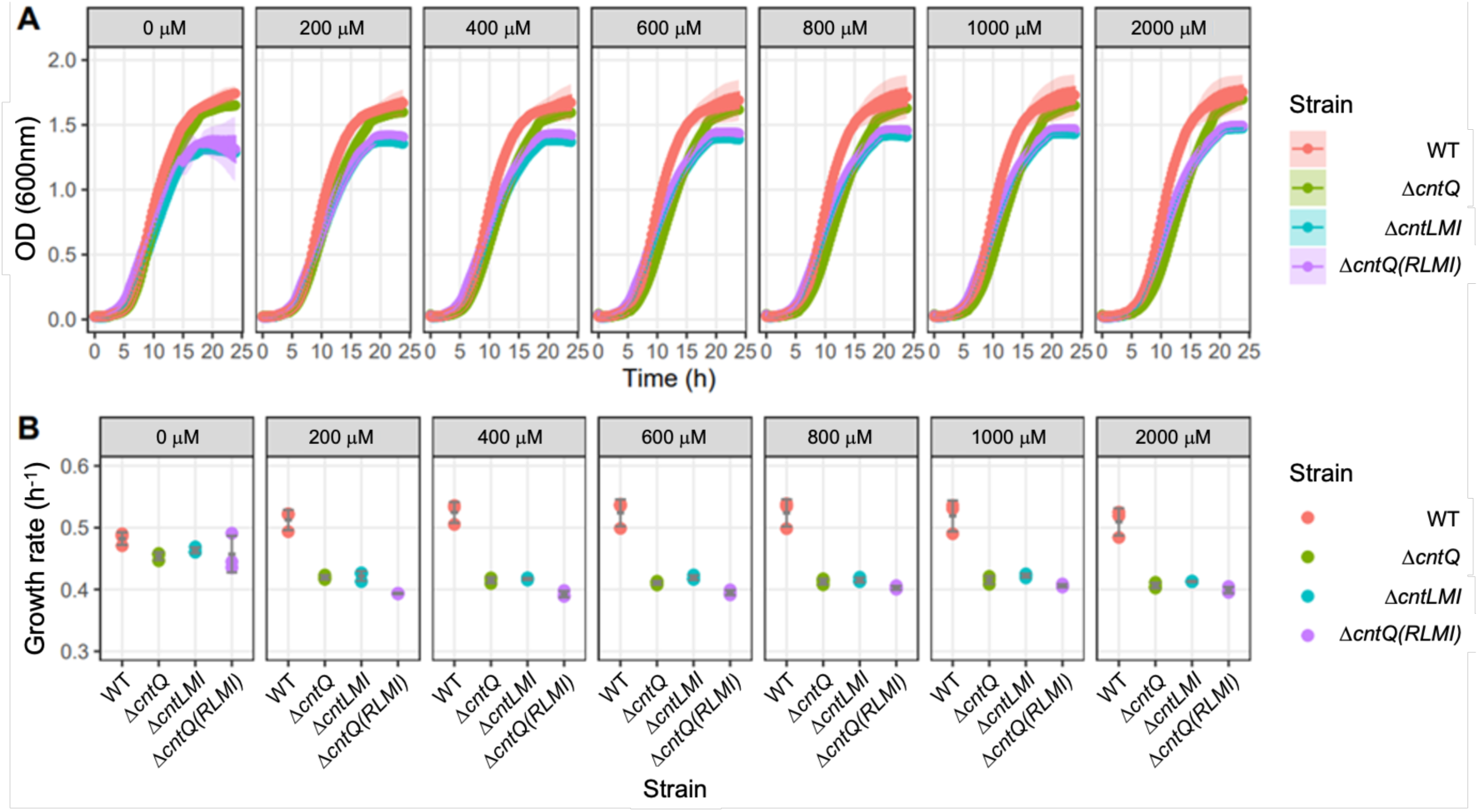
Growth curves (A) and inferred growth rates (B) of *Y. pseudotuberculosis* cultivated in TMH without or with increasing concentration of EDTA. In A, the data represent the means and the confidence intervals (95%) calculated from three biological replicates. In B, the data from these three biological replicates are shown as individual points associated with their mean and standard deviation (in grey).

**Supplementary Table 1.**
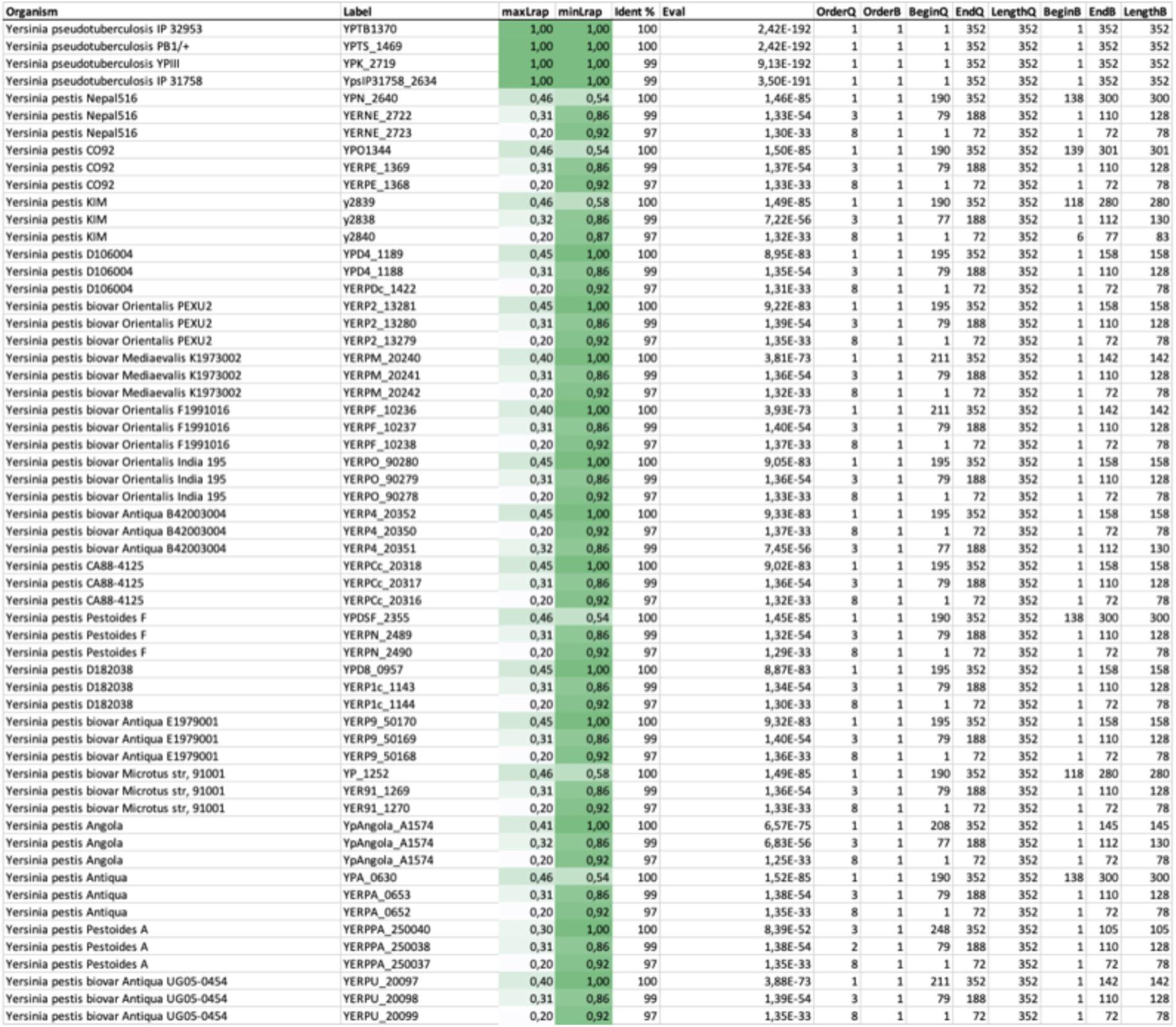
Pseudogenization of the *cntQ* permease is conserved in *Y. pestis* strains. List of orfs predicted in sequenced *Y. pseudotuberculosis* and *Y. pestis* strains at the *cntQ* position. maxLrap and minLrap denote respectively the length (expressed as a percentage of the gene) that is conserved, taking YPTB1370 as a reference (Query). BeginQ and EndQ refer to portions of sequence that is conserved on YPTB1370, taking YPTB1370 (Query) as a reference. BeginB and EndB refer to portions of sequence that is conserved in a given sequence (Bait), using YPTB1370 as a reference.

**Supplementary Table 2.**
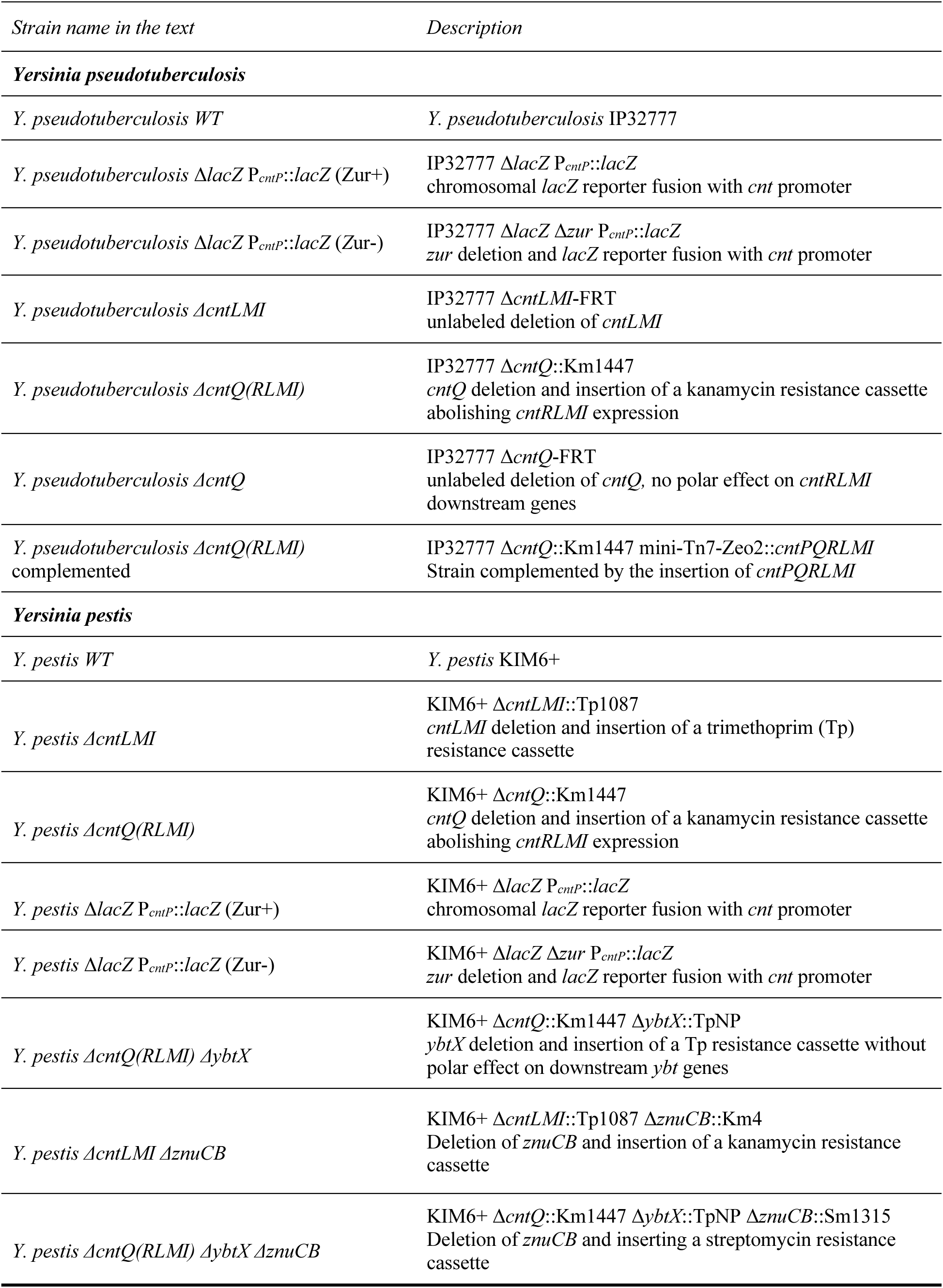
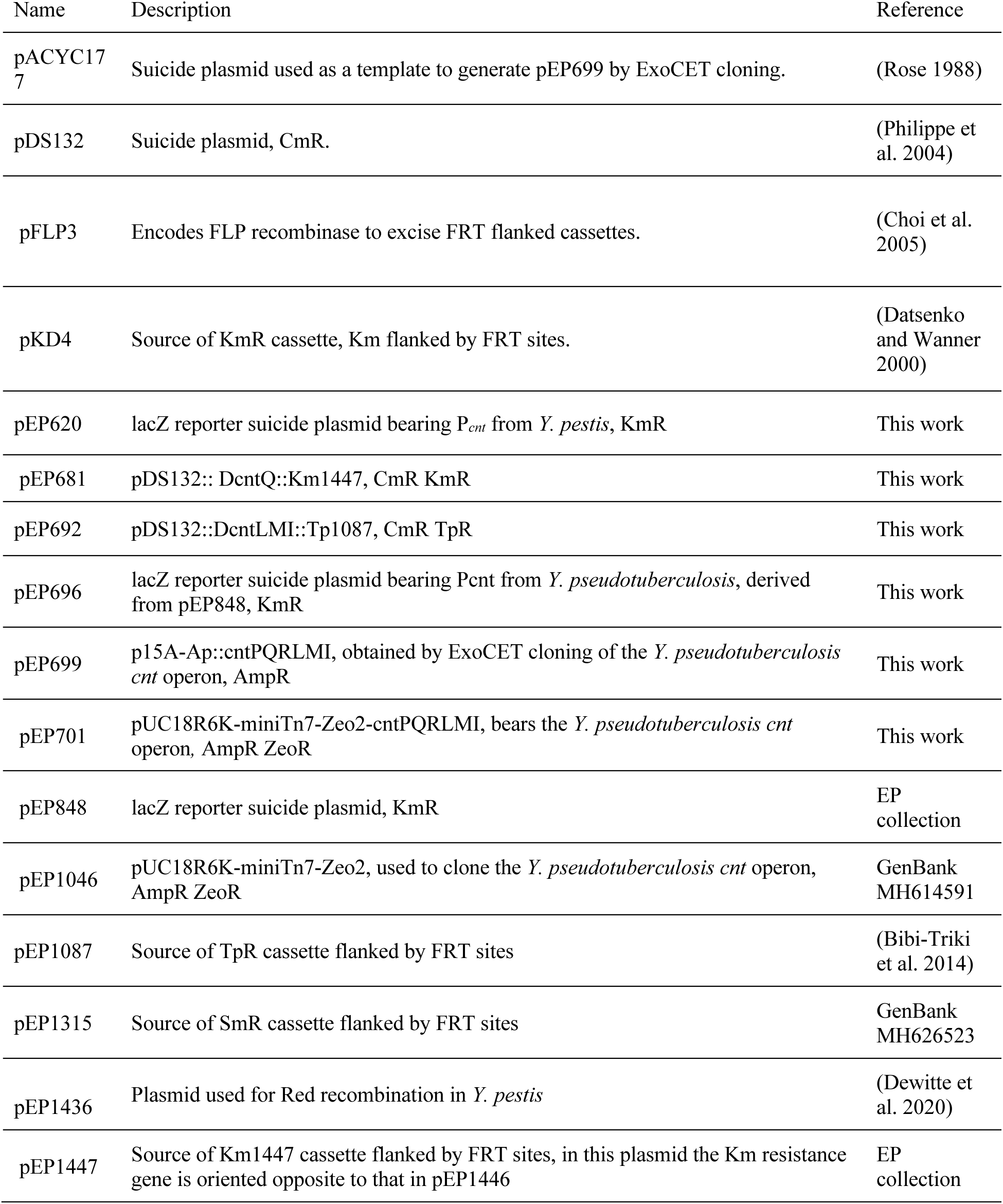
Bacterial strains and plasmids used in this study.

**Supplementary Table 3.**
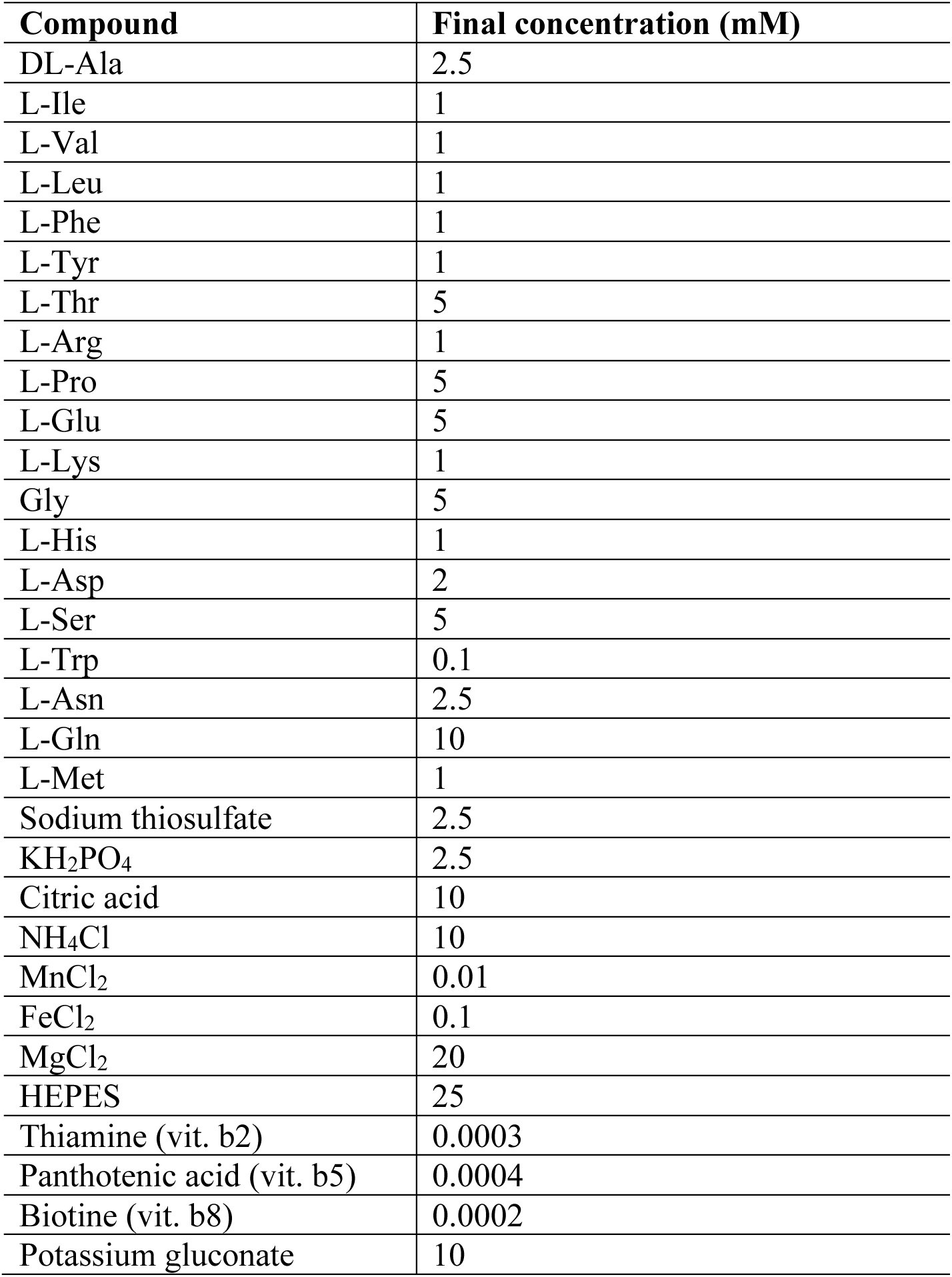
Composition of the TMH medium.

**Supplementary Table 4.** Metal concentrations in TMH medium. Metal concentration as determined by ICP-AES (Bdl: below detection level).

| **Metal** | **Concentration (μM)** |
| --- | --- |
| Mn | 6.3 |
| Fe | 62.5 |
| Zn | Bdl |
| Cu | Bdl |
| Ni | Bdl |
| Co | Bdl |

